# Forces driving cell sorting in *Hydra*

**DOI:** 10.1101/142976

**Authors:** Olivier Cochet-Escartin, Tiffany T. Locke, Winnie H. Shi, Robert E. Steele, Eva-Maria S. Collins

## Abstract

Cell sorting, whereby a heterogeneous cell mixture organizes into distinct tissues, is a fundamental patterning process in development. So far, most studies of cell sorting have relied either on 2-dimensional cellular aggregates, *in vitro* situations that do not have a direct counterpart *in vivo,* or were focused on the properties of single cells. Here, we report the first multiscale experimental study on 3-dimensional regenerating *Hydra* aggregates, capable of reforming a full animal. By quantifying the kinematics of single cell and whole aggregate behaviors, we show that no differences in cell motility exist among cell types and that sorting dynamics follow a power law. Moreover, we measure the physical properties of separated tissues and determine their viscosities and surface tensions. Based on our experimental results and numerical simulations, we conclude that tissue interfacial tensions are sufficient to explain *Hydra* cell sorting. Doing so, we illustrate D’Arcy Thompson’s central idea that biological organization can be understood through physical principles, an idea which is currently re-shaping the field of developmental biology.

**Summary statement:** *Hydra* regenerates after dissociation into single cells. We show how physical mechanisms can explain the first step of regeneration, whereby ectodermal and endodermal cells sort out to form distinct tissue layers.

## Introduction

How a pattern emerges from an initially near-uniform cell population is a question that has long fascinated biologists and physicists alike, in particular d’Arcy Thompson. In his influential 1917 book *On Growth and Form* (Thompson, 1992), Thompson emphasized the fact that, when one is faced with such a complex phenomenon as the form of a living organism, there can be more than one explanation, depending on the level of understanding one aims to achieve (molecular, cellular, organismal). Although evolutionary and molecular processes play key roles in morphogenesis, Thompson insisted on the importance of studying this question from a purely physical perspective: “My sole purpose here is to correlate with mathematical statement and physical law certain of the simpler outward phenomena of organic growth and structure or form […]. But I would not for the world be thought to believe that this is the only story which Life and her Children have to tell” (Thompson, 1992).

One of the simplest and best studied examples of pattern formation in which this approach has been fruitful is the separation of two cell populations which have been mixed to yield a heterogeneous cell suspension, in a process called cell sorting. Since the dynamics of cell sorting resemble the breaking up of an emulsion of different liquids, physically-based mechanisms have long been suggested to explain this process (reviewed in (Foty and Steinberg, 2013)). From a physics perspective, cell populations (tissues) are active, complex fluids. They are active because cell motility is driven by ATP consumption and not by thermal energy. They are complex because they exhibit elastic solid-like behavior on short timescales and viscous liquid-like behavior on long timescales (Forgacs et al., 1998). Examples of viscous liquid-type behaviors are rounding of tissue pieces and fusion of tissues upon contact (Schötz et al., 2008). In liquids, both of these processes are driven by surface tension. Accordingly, the “Differential Adhesion Hypothesis” (DAH) proposed that cell sorting is a direct consequence of differences in tissue surface and interfacial tensions, similar to the breaking up of an emulsion (Steinberg, 1970). When cells from two tissue types are mixed and able to interact via cell adhesion, they will sort according to their respective tissue surface tensions, whereby the tissue with lower surface tension engulfs the tissue with the higher surface tension (Foty et al., 1996). Molecularly, tissue surface tensions were originally attributed to differences in adhesion alone (Duguay et al., 2003; Foty and Steinberg, 2005), but have since been shown to arise from an interplay between cell adhesion and cortical tension (Manning et al., 2010).

In the theoretical work of Glazier and Graner (1992), a Cellular Potts Model (CPM) was created to simulate the behavior of single cells during cell sorting (Graner and Glazier, 1992). They demonstrated that differences in interfacial energies were sufficient to drive the spontaneous sorting of two cell populations (Glazier and Graner, 1993; Graner, 1993). Since then, this and other models have been refined to include other mechanisms such as coherent motion (Belmonte et al., 2008), biochemical dynamics of adhesion molecules (Zhang et al., 2011), or chemotaxis (Vasiev and Weijer, 1999). In all cases, differences in tissue surface tensions drive sorting, but the resulting dynamics are modified by these additional mechanisms. Furthermore, others have shown that sorting can occur in the absence of differences in tissue surface tension. Such models mostly rely on asymmetries of cell motility to explain sorting, either from intrinsic differences between cell types (Beatrici and Brunnet, 2011) or from differences in a cell’s immediate surroundings (Strandkvist et al., 2014). However, there is disagreement on the rules regulating engulfment for two cell types with different locomotion properties. Jones and colleagues (1989) found that chick tissues sorted such that the fastest moving tissue ended up on the inside of a mixed cellular aggregate (Jones et al., 1989). In contrast, theoretical work showed that when full sorting occurred, faster cells surrounded slower ones and formed streams around them (Beatrici and Brunnet, 2011).

The freshwater cnidarian *Hydra* has frequently been the model of choice for studies of cell sorting. *Hydra* is anatomically simple with radial symmetry and two epithelial tissue layers, ectoderm and endoderm. *Hydra* can be dissociated into individual cells which, when aggregated, can autonomously regenerate whole animals (Gierer et al., 1972). Cell sorting into a sphere-within-a-sphere configuration, with an inner endoderm and outer ectoderm, is the first step in this regeneration process and a necessity for subsequent developmental milestones - the formation of a hollow bilayered sphere, symmetry breaking, and axis formation (Gierer et al., 1972). Therefore, cell sorting in *Hydra* is experimentally accessible to quantitative studies while remaining physiologically relevant.

Differential surface tension was hypothesized to be key to *Hydra* cell sorting. Support for this view came from centrifugation experiments, which showed that under similar centripetal forces, endodermal epithelial cells formed larger aggregates than ectodermal epithelial cells, indicating that endoderm has a higher tissue surface tension than ectoderm, in agreement with the DAH (Technau and Holstein, 1992).

A direct measurement of adhesion strength of epithelial cell pairs using optical traps (Sato-Maeda et al., 1994) found that adhesion between endodermal epithelial cells is stronger than adhesion between ectodermal epithelial cells, in agreement with a DAH-driven sorting process. However, the authors found that heterotypic cell-cell interactions were the weakest of all, in disagreement with the DAH framework which requires that the heterotypic interaction strength be intermediate between the strongest (endo/endo) and weakest (ecto/ecto) interactions. One possible explanation for this discrepancy is time-dependent changes in cell-cell interaction strengths. This idea was confirmed by more recent work which found that cell sorting of *Hydra* aggregates may have two phases: a short initial phase in which homotypic cell interactions dominate and ectodermal-endodermal interaction does not occur (Hobmayer et al., 2001), and a second longer phase, in which ectoderm displays a higher affinity for endoderm than for itself. Since the aggregates investigated in this study were small and non-viable, whether the existence of a short initial phase is relevant for the sorting of large aggregates capable of regenerating into full animals (10^3^-10^4^ cells) is unknown.

To test whether tissue surface tensions and adhesion differences between ectoderm and endoderm were sufficient to explain sorting or whether other parameters had to be considered, other studies investigated single cell behaviors. For example, Takaku et al. (2005), studied the behavior of isolated ecto- or endodermal cells when put in contact with a tissue sphere. They found that a single ectodermal cell in contact with an endodermal aggregate does not migrate into the aggregate, whereas a single endodermal cell in contact with an ectodermal aggregate does migrate to the interior (Takaku et al., 2005). They interpreted this finding as indicative of differences in cell motility, although this behavior is also expected from the DAH. Additional experiments were performed that seemed to reveal such differences in motility. For example, they showed that epiboly, the process by which an ectoderm aggregate spontaneously engulfs an endoderm aggregate upon contact, depends on the motility of endodermal but not of ectodermal cells (Takaku et al., 2005). Furthermore, they found that in clusters of 4 cells (2 endodermal and 2 ectodermal), ectodermal homotypic adhesion was more stable than endodermal homotypic adhesion, seemingly in contradiction to the DAH (Takaku et al., 2005). However, since the stability of an adhesion depends not only on its strength but also on the activity of the cells forming it, and endodermal cells were observed to be more actively motile, it is possible that endoderm-endoderm adhesions only appeared weaker.

Other studies have focused on quantifying cell motility during cell sorting and within homotypic tissues without addressing the underlying mechanism driving sorting. Rieu et al. quantified the motility of endodermal cells in aggregates of different compositions (Rieu et al., 1998; Rieu et al., 2000): pure endoderm, pure ectoderm, and evenly mixed. Overall, their results show that cells move in a mostly random fashion and are most mobile in a purely ectodermal environment. This result agrees with the finding in (Takaku et al., 2005), but is not conclusive regarding differential cell motility. Moreover, this result is also expected from the DAH, since an ectoderm aggregate would make for a less cohesive environment in which a single cell can move more freely. In summary, both theoretically and experimentally, the existing data on *Hydra* cell sorting are insufficient to delineate whether sorting is driven by one of the two proposed classes of models. To achieve a definitive explanation, more quantitative data on the behavior of individual cells and both tissues in a physiologically relevant 3-dimensional setting, as well as direct measurements of tissue surface tensions, are needed.

Here, we take advantage of transgenic *Hydra* (Wittlieb et al., 2006), to revisit this long standing question about the mechanisms of cell sorting through a multiscale approach. We first focus on 3-dimensional mixed aggregates - large enough to regenerate into polyps - and present quantitative data on the dynamics of sorting. To determine whether differential surface tensions can drive sorting, we performed rheological measurements of both tissues’ mechanical properties, in particular of their surface tensions. Next, we mapped single cell trajectories within mixed aggregates to address whether the two cell types possess intrinsically different motile properties. Finally, we developed numerical simulations using the 3D Cellular Potts Model (CPM) based on our experimental conditions, and compared the *in silico* results to our experimental data. We find that differences in tissue surface tensions are indeed sufficient to reproduce all of our experimental data.

In summary, our work explains the physical mechanism by which *Hydra* cell aggregates sort into tissues. Surface tension-driven demixing of ectodermal and endodermal epithelial cells is the first critical step in the regeneration of the whole animal after complete dissociation. Thus, by explaining how cell sorting works in *Hydra* aggregates, we are one step closer to a complete understanding of biological pattern formation.

Our work illustrates by one example the core idea of D’Arcy Thompson’s book. While cell sorting can be understood in terms of molecular mechanisms (motility, adhesion, cortical tension), we show that it can also be understood, at a coarse-grained level, using the physics of liquids without detailed knowledge of the underlying molecular machinery.

## Results

A natural starting point for distinguishing between the most prominent explanations of cell sorting in *Hydra,* i.e. differential tissue surface tension versus differential motility, is to perform measurements on the cell sorting dynamics. Second, as the ingredients in these models are either based on tissue rheological properties or single cell motility, we quantitatively assessed these two aspects.

## Dynamics of cell sorting in *Hydra* aggregates

All models of *Hydra* cell sorting predict that the two initially mixed cell populations spontaneously separate, with ectoderm engulfing endoderm. The dynamics of sorting, however, depend on the model ingredients and their analysis could therefore possibly enable a distinction between the different mechanisms. To test this, we prepared cellular aggregates from transgenic *Hydra* in which the two epithelial layers express different fluorescent proteins. Initially, aggregates showed a random mixture of both cell types (Fig. 1A) and were disc-shaped, because cells are re-aggregated via centrifugation (see Methods). Over the course of 4-10 hours, the two cell populations spontaneously separated and the disc-shaped aggregate rounded up into a solid sphere (Movie 1). The ectodermal cells moved toward the outside while the endodermal cells moved toward the center of the aggregate, leading to a sphere-within-a sphere configuration. Once sorting was complete, the aggregates ejected excess cells as they transitioned into a hollow bilayer epithelial sphere. The bilayer sphere eventually broke symmetry and regenerated into an adult polyp (Fig. S1). *Hydra* cell aggregates are therefore a true *in vivo* system, despite their apparent simplicity. Indeed, regeneration from aggregates occurs even in epithelial *Hydra* which have been reduced to ectoderm and endoderm through removal of the interstitial cell lineage (Marcum and Campbell, 1978). Interstitial stem cells and their progeny thus do not significantly alter cell sorting and the system can be treated as a two-component mixture. The other potentially important player in cell sorting could be the extra-cellular matrix (ECM), which separates the two epithelial tissues in the intact animal. However, using antibody staining we verified that laminin, a major component of *Hydra* ECM, was undetectable during sorting (Fig. S2), in agreement with previous reports (Seybold et al., 2016).

**Figure 1.**
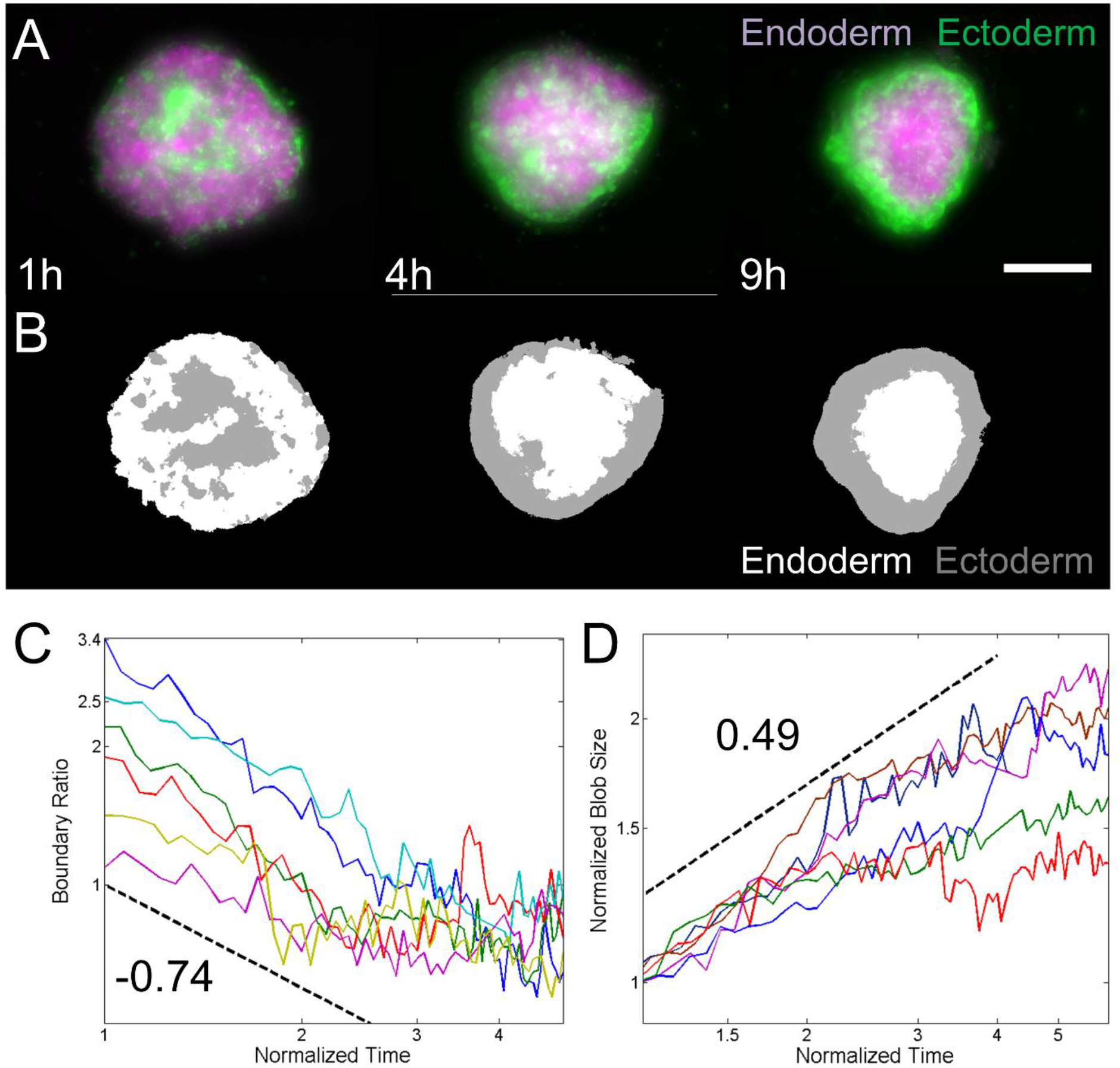
Dynamics of cell sorting. *A) Representative still images of sorting of Hydra watermelon aggregates capable of regeneration. The reduction in radius is a signature of the aggregate rounding up in three dimensions. B) Automated image analysis of the images in Panel A determining the position of both tissues. Scale bar: 200 μm. C) Log-log plot of boundary ratio as a function of normalized time for six representative experiments. The dashed black line shows the behavior of a power law with exponent -0.74. D) Log-log plot of normalized blob size as a function of normalized time for five representative experiments. The dashed black line shows the behavior of a power law with exponent 0.49. In panels C and D, the long term behavior shows saturation of these measurements and therefore deviation from a power law behavior.*

To allow for a direct comparison of our experimental results with existing theoretical models and predictions, we focused on measuring quantities that are commonly used in the field. One such quantity is the sorting index, which measures the average fraction of neighboring cells that are of the opposite type. In an evenly mixed aggregate, the initial value of the sorting index is 0.5. It decreases as sorting proceeds and saturates at a value that depends on the system size. The sorting index is difficult to measure experimentally as it requires knowledge of the position, neighbors, and identity of all cells within an aggregate. However, the sorting index is directly proportional to the length of the boundary (see Methods) between the tissues and therefore follows the same functional form. With the exception of a more complex model taking into account the biochemical dynamics of adhesion proteins (Zhang et al., 2011), models based on either DAH or differences in motility have both found that this decrease follows a power law. The exponents, however, vary depending on the details of the model such as the ratio of cells from both types (Nakajima and Ishihara, 2011) or the differences in motility built into the model (Strandkvist et al., 2014).

Using automated image analysis (Fig. 1B), we measured the length of the boundary between endoderm and ectoderm as a function of time (see Methods). We found the boundary length decrease to follow a power law (Fig. 1C) with exponent -0.74+-0.24 (mean+-STD n=17, Fig. S2). This exponent is significantly higher than those reported in various theoretical works which ranged from -0.025 to -1/3 (Beatrici and Brunnet, 2011; Belmonte et al., 2008; Nakajima and Ishihara, 2011; Strandkvist et al., 2014), implying that the observed sorting is faster than previously suggested. We present a more detailed analysis and explanation of this result in the discussion section.

Another quantity used in the field is the typical blob (cluster) size of both tissues as a function of time. The definition of the typical blob size varies from one study to the next. Since this measure is also commonly used in the study of phase separation through spinodal decomposition (Fan et al., 2016), we chose here to use the same definition (see Methods). This allowed us to directly compare our dynamics to a purely physical situation. The blob size is linked to the total sorting time, as sorting is complete once the typical blob size reaches a value comparable to the system size.

We found that the typical blob size increases as a power law (Fig. 1D), with an exponent of 0.49+-0.24 (mean+-STD, n=17, Fig. S3). Again, this implies faster sorting than previously reported (Belmonte et al., 2008; Glazier and Graner, 1993; Nakajima and Ishihara, 2011). Of note, scaling rules imply that blob size and boundary ratio should have equal exponents of opposite sign. Our mean values are quite different (0.74 versus 0.49) but still within experimental uncertainties of each other. The real exponent is likely intermediate between these values.

In summary, studying cell sorting dynamics is instructive, but was insufficient to distinguish between different sorting mechanisms. This is due to the fact that our experimentally determined exponents differ from published values; we therefore cannot draw conclusions on the sorting mechanism from these experiments alone.

However, because the different models for cell sorting also make assumptions and predictions regarding the properties of each tissue separately and/or on the behavior of single cells within aggregates, we performed experiments at these scales to further probe the possible mechanisms explaining cell sorting.

## Physical behavior of separated tissues

The DAH is based on the assumption that each tissue behaves like a liquid on long time scales. Sorting is then driven by the effective interfacial and surface tensions of the two tissues. Cells, like liquid molecules, have a lower contact free energy with each other than with the medium. Cellular rearrangements thus tend to minimize the tissue’s contact area with this medium, which is similar to the effect of surface tension in liquids (Forgacs et al., 1998; Foty et al., 1994).

For full sorting and engulfment to occur, these rheological quantities need to obey the following inequality (Foty et al., 1996):

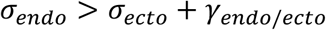

Where *σ* represent the tissue’s surface tensions and *γ*_endo/ecto_ their interfacial tension.

DAH thus predicts that each tissue flows on long time scales and that the effective surface tension of endoderm is higher than that of ectoderm. Because these questions have to be addressed for each tissue separately, this requires their physical separation. To do so, we adapted previously published protocols (Kishimoto et al., 1996) (see Methods) to chemically dissolve the ECM which leads to the physical separation of the tissues.

By this separation, we obtained tissue pieces containing only one of the two epithelial cell types, and then observed their long time behaviors. Tissue pieces rounded up (Movie 2) and fused (Movie 3) on time scales of minutes to a few hours, thus demonstrating liquid behavior. This justifies the usage of concepts like surface tension and viscosity. To determine both of these quantities, we used micro-aspiration experiments (Fig. 2A-C, Movie 4), in which tissue pieces are aspirated into a micro-pipet using negative pressure. The pressure needed to initiate aspiration is directly related to the surface tension of the tissue of interest (Guevorkian et al., 2010) according to:

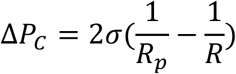

Where Δ*P_C_* is the critical pressure required to trigger aspiration, *σ* is the surface tension, *R_p_* is the radius of the micro-pipet and *R* that of the tissue piece.

**Figure 2.**
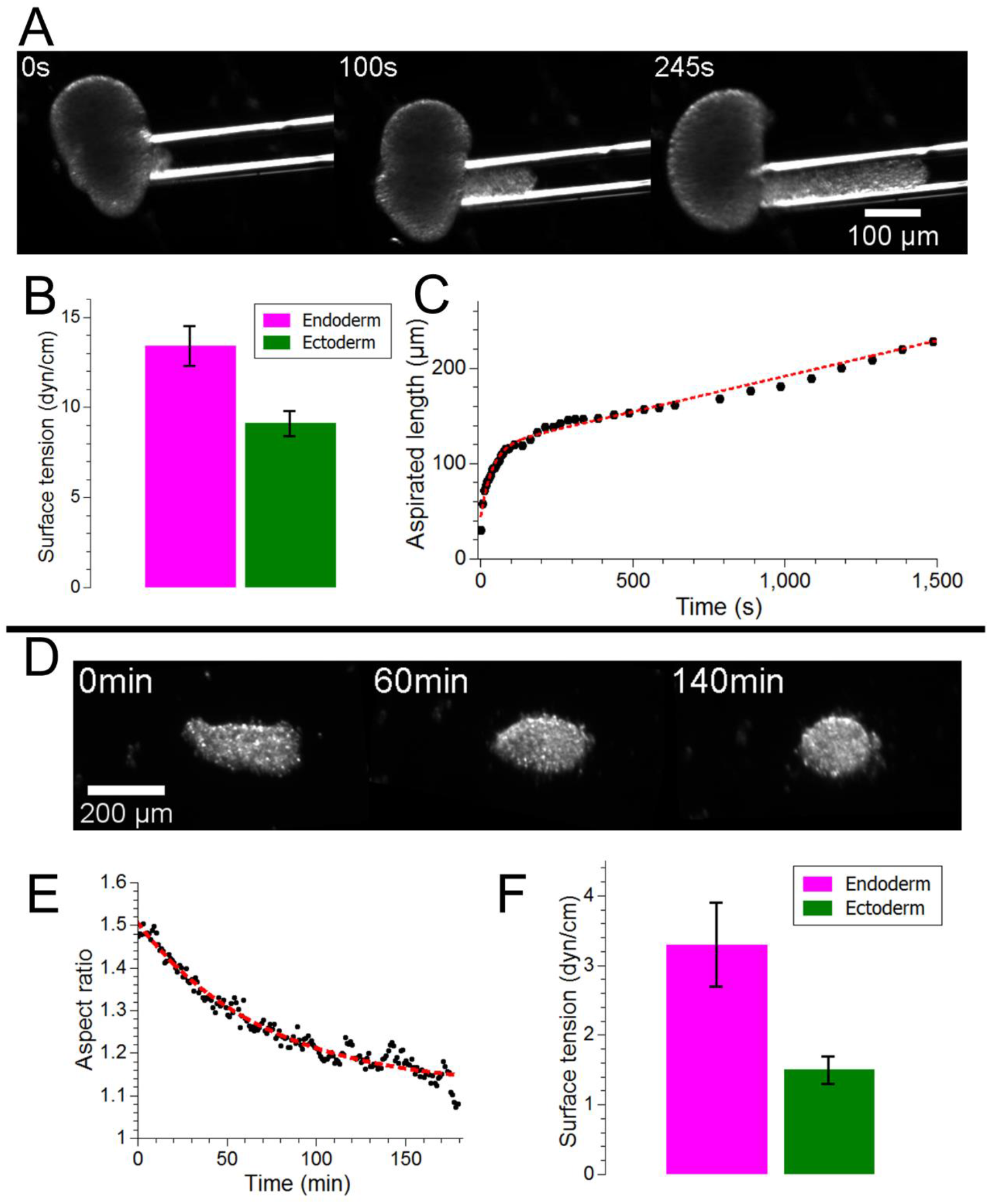
Rheology of individual tissues. *A) Still sequence of micro-aspiration experiment performed on an endoderm tissue piece. B) Quantification of surface tensions from micro-aspiration experiments, the bars shows mean ± SEM, n=13 and 14 for endoderm and ectoderm, respectively. C) Sample quantification of aspirated length of an endoderm piece as a function of time showing a short, visco-elastic phase used to estimate viscosity and a long linear phase. D) Still sequences of rounding up experiment on an ectoderm tissue piece. E) Quantification of aspect ratio as a function of time of the experiment shown in D), the dashed red line represents an exponential fit to the data in black. F) Quantification of surface tensions from rounding up experiments, the bars show mean ± SEM, n=17 and 15 for endoderm and ectoderm, respectively.*

We estimated surface tensions this way and found the surface tension of endoderm to be higher than that of ectoderm (Fig. 2B, Table. S1), in agreement with the DAH (13.4 ± 1.1 dyn/cm and 9.1 ± 0.7 dyn/cm, respectively, mean ± SEM, n=13 and 14). In addition, the first phase of aspiration (Fig. 2C) is dominated by the visco-elastic response of the tissues leading to an exponential relaxation. The characteristic time of this relaxation *τ* is given by (Guevorkian et al., 2010):

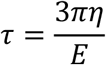

Where *E* and *η* are the elastic modulus and viscosities, respectively. Using previous measurements of the elastic moduli with parallel plate compression (Carter et al., 2016), we found the viscosities of endoderm and ectoderm to be 3.7 ± 0.7 10^4^ Pa.s and 4.8 ± 0.6 10^4^ Pa.s, respectively (mean ± SEM, n=10 and 9, Table. S1), similar to measurements performed on other cell aggregates including from different chicken embryonic cells (Forgacs et al., 1998) and mouse sarcoma (Guevorkian et al., 2010; Marmottant et al., 2009).

The viscosity estimates can further be used to independently determine tissue surface tensions from rounding up experiments, because this behavior is driven by surface tension and slowed down by viscosity. Similar to what has been reported for other cellular aggregates (Mombach et al., 2005), the dynamics of a tissue piece rounding up, measured as the decrease of the piece’s aspect ratio over time, was exponential (Fig. 2E). The characteristic time *τ* of this exponential relaxation is given by:

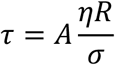

Here *R* is the radius of the tissue piece and *A* is a numerical pre-factor which depends on geometry. This numerical pre-factor has been estimated to be on the order of 0.95 in different circumstances (Gordon et al., 1972), but is unknown in our case. Since we are primarily interested in the relative differences between the surface tensions of both tissues, knowledge of this pre-factor is not crucial. Using the viscosity measurements obtained by microaspiration, we found a higher surface tension for endoderm when compared to ectoderm (3.3±0.6 and 1.5±0.2 dyn/cm, respectively, mean ± SEM, n=17 and 15) (Fig. 2F, Table S1). We attribute the difference of these absolute values from the micro-aspiration results to the undetermined prefactor in rounding. Importantly, however, the relative differences in surface tensions obtained through both methods are of the same order.

In principle, fusion experiments could be similarly used to estimate surface tension.

However, while endoderm tissue pieces readily fused (Movie 5), ectoderm pieces fused rarely and only if freshly cut surfaces were directly brought into contact (Movie 5). We attribute this difference in behavior to the polarity of ectoderm pieces, which was previously reported (Takaku et al., 2005). Because of polarization, cells in contact with the outside may be non-adhesive to the outside and thus unable to fuse with a neighboring piece. This lack of boundary cell-cell interaction would have no effect on rounding up and micro-aspiration, and little effect on cell sorting, since polarization would only happen once ectodermal cells reach the aggregate boundary. Thus, while polarization can be neglected in these other experiments, it dominates ectoderm behavior during fusion and thus complicates estimates of tissue surface tensions from fusion experiments.

Finally, we repeated a qualitative experiment, previously performed by Technau and Holstein (Technau and Holstein, 1992). We showed that under similar aggregating forces and at similar cell densities, endodermal cells made larger aggregates than ectodermal ones, a signature of their highest cohesiveness (Fig. S4), in agreement with experiments performed at the single cell level (Sato-Maeda et al., 1994).

Overall, we demonstrated that both tissues show liquid-like behaviors on long time scales (rounding up, flowing) and that the endoderm has a higher surface tension than the ectoderm.

The difference is high enough to explain cell sorting (Foty et al., 1996). Our estimates of tissue viscosities and surface tensions are in good agreement with previously published values on aggregates of embryonic tissues from chicken (Forgacs et al., 1998; Foty et al., 1994; Foty et al., 1996) or zebrafish (Schötz et al., 2008) which were all on the order of 10^4^-10^5^ Pa.s for viscosities and 1-30 dyn/cm for surface tensions. Together these results demonstrate that differential surface tension plays an important role during cell sorting in *Hydra* aggregates.

## Single cell dynamics during sorting

Our results show that differences in surface tension can drive cell sorting but they do not exclude the involvement of other mechanisms such as differences in cell motility. To evaluate whether differential cell motility played a role in cell sorting, we tracked individual cells during the sorting process. To achieve single cell tracking within the aggregates, we prepared aggregates in which 5% of the cells were transgenic, expressing a fluorescent protein. These aggregates were analyzed using three dimensional 2-photon timelapse imaging. From the resulting videos, we reconstructed single cell trajectories for both tissues, in the first 4-6 h of sorting (Fig. 3A). We found cell speeds to be on the order of 50 μm/h, constant throughout this time window, and comparable for both cell types (Fig. 3B). The mean square displacements (MSDs) of both cell types were weakly super-diffusive (power law with exponent of 1.3-1.4) and, again, similar (Fig. 3C). This indicates that cell motion was mostly random and thus that directed cell motility doesn’t play a role in cell sorting. This is further demonstrated by the fact that cell directionality was also similar for both cell types, despite their differing final positions (inside versus outside) (Fig. 3D). Quantitatively, the MSDs yielded diffusion coefficients on the order of a few hundred microns squared per hour, the same order of magnitude as was reported for two dimensional *Hydra* aggregates (Rieu et al., 1998).Finally, we found no differences in speed distributions and velocity auto-correlation functions between the two cell types (Fig. S5).

**Figure 3.**
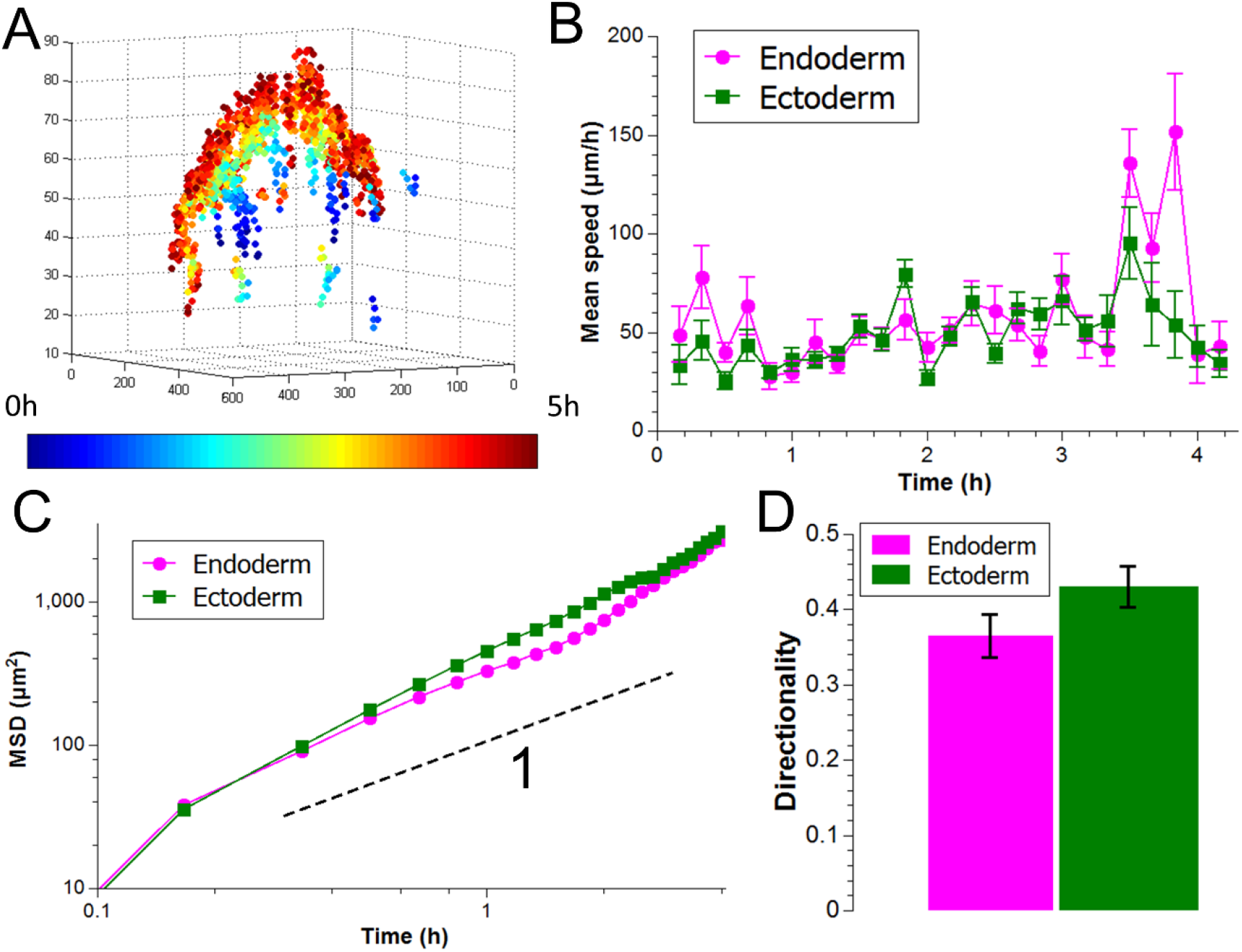
Single cell dynamics. *A) Reconstructed cell tracks from 2-photon imaging color coded by time. Axes are in microns. B) Quantification of mean speeds of both tissues from one representative experiment. C) Log-log plot of mean square displacements in the same experiment. The black dashed line shows the behavior of a power law with an exponent of 1, i.e. purely diffusive motion. This shows that cell motion is slightly super-diffusive. A linear fit yields diffusion constants of 564±63 μm^2^/h and 657±23 μm^2^/h (best fit ±95% confidence interval) for endoderm and ectoderm, respectively. D) Quantification of directionality from the same representative experiment. Directionality is averaged over all traceable cells and the bars represent mean±standard deviation (n=20 and 17 for endoderm and ectoderm, respectively)*.

Of note, we observed a general vertical trend in the displacement of the cells which is explained by the fact that aggregates start in a mostly flat state and round up as they sort. To test if our results were dominated by the global motion of the aggregate, we performed experiments where all nuclei were stained to correct for this global motion and obtained similar results (Fig. S6). This shows that, although important, the global motion of the aggregate does not impact our conclusions. The main difference was that center-of-mass motion-corrected MSDs were basically linear (Fig. S6), implying that the coherent component of the cell motion is due to rounding up and not cell sorting, again in agreement with experiments performed in two dimensions (Rieu et al., 2000).

In summary, our data at the single cell level do not reveal any intrinsic motility differences between the two tissue types. This implies that differential motility does not play a role in cell sorting in *Hydra* aggregates.

## DAH-based numerical simulations of cell sorting

Since our experiments showed that differential surface tension governs cell sorting without differential motility playing a significant role, we used numerical simulations to probe the effects of both mechanisms and test their ability to reproduce our experimental data. We applied a cellular Potts model to simulate cell sorting using the freely available CompuCell 3d software (http://www.compucell3d.org/)(Swat et al., 2012). The simulations are based on differential adhesive forces between pairs of cells depending on their identities. Individual cells tend to keep their volumes (finite compressibility) and their surface area (finite deformations). To mimic our experiments as closely as possible, simulations were run in 3 dimensions using different numbers of cells. Initially the aggregates were in a flat configuration with a thickness of 3 cells and the long sides were varied from 7.5 to 35 cells (Fig. 4A) leading to a total number of cells ranging from a hundred to a few thousand. Although the largest simulated aggregates were smaller than the largest aggregates in the experiments (on the order of 10^4^ cells), they were large enough to model aggregates capable of regeneration. In addition, we tuned the adhesion parameters to obtain a surface tension twice as high for the endoderm as for the ectoderm (see Methods), the same order of magnitude found in our experimental measurements.

As expected, we found that these features were sufficient to drive both cell sorting (Fig. 4A, Movie 6) and the rounding up of the aggregate observed in experiments (Fig. 1A).

Regarding single cell dynamics, both cell types showed similar motility behaviors as shown by their respective speeds, MSDs, or directionality (Fig. 4B-D). We found the MSDs to be slightly super-diffusive (exponent of 1.2-1.3), in agreement with our experimental results (Fig. 3C). To quantitatively reconcile length scales between simulations and experiments, we used typical cell sizes (4 pixels in simulations and 20 microns in experiments). In simulations, as in experiments, we found that individual cell speeds were constant and similar for both tissues. We thus decided to equalize the speeds that led to each simulation step to be on the order of 10s. This led to sorting times ranging from 1h to 10h, in agreement with experimental observations (Fig. S7).

**Figure 4.**
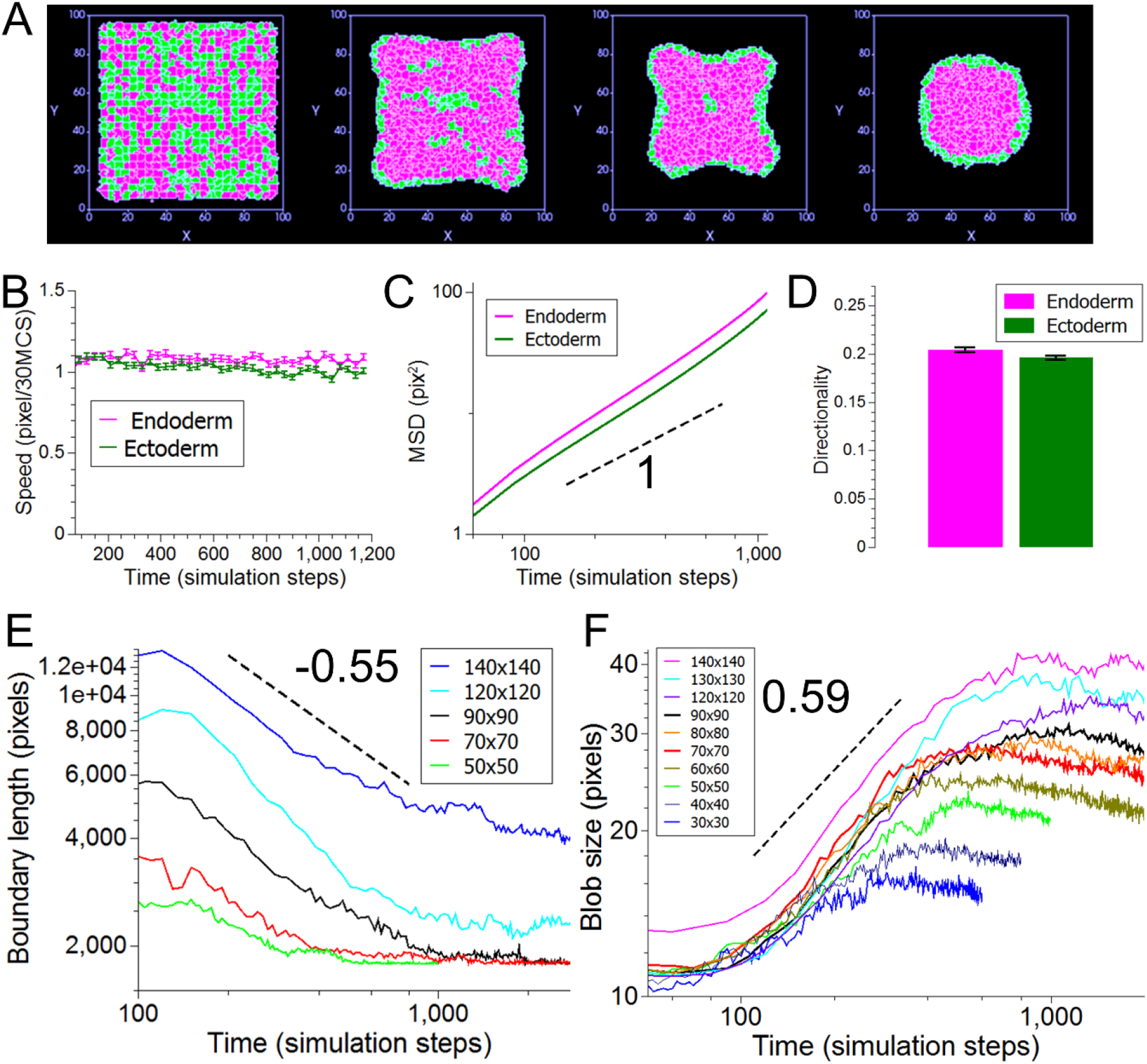
DAH-based numerical simulations. *A) Still sequence of the middle slice in a simulation. Endodermal cells are shown in magenta and ectodermal cells are shown in green. Stills are shown at 0, 300, 800, and 2000 simulation steps. B) Measurement of speeds over time for a representative simulation at the largest size. C) Log-log plot of the mean square displacement of the same simulation as in B). The black dashed line shows the behavior of a power law with an exponent of 1 or diffusive behavior. A linear fit yields diffusion constants of 0.084±0.004 pixel^2^/simulation steps and 0.07±0.004 pixel^2^/simulation step (best fit±95% confidence interval). D) Quantification of directionality from the same representative simulation. Directionality is averaged over all tractable cells and the bars represent mean±standard deviation (n=1837 for both tissues) E) Log-log plot of boundary length as a function of time for five different initial sizes. Each plot represents the mean of three independent simulations. The dashed black line shows the behavior of a power law with exponent -0.55. F) Log-log plot of blob size as a function of time for ten different initial sizes. Each plot represents the mean of three independent simulations. The dashed black line shows the behavior of a power law with exponent 0.59*.

For aggregate-scale dynamics in the simulations we found that both the length of the boundary between the two tissues (Fig. 4E) and the typical blob size (Fig. 4F) followed power laws. For larger aggregates, these exponents were independent of size (Fig. S8). The boundary length decreased as the power -0.55±0.11 of time while the blob size increased as the power 0.59±0.03 (mean ± standard deviation, n=21 simulations at seven different initial sizes). These values are both within error bars of what we obtained in the experiments.

Finally, our model also correctly reproduced the fluid-like behavior of separated tissues as shown by simulations of fusion (Movie 7) and engulfment (Movie 8). Taken together, these results demonstrate that a numerical simulation based solely on DAH, reproducing the geometry of our experiments and with model parameters partly coming from experimental measurements (see Methods) was sufficient to reproduce the data we gathered at different scales. Therefore, we conclude that DAH is sufficient to explain cell sorting in three-dimensional *Hydra* aggregates.

It is possible, however, that differential cell motility acts in addition to differential interfacial tensions and speeds up sorting. To test the effect of adding differential motility, we ran simulations including both mechanisms by separately tuning the effective temperatures of both tissues. Clearly, cellular processes are not driven by thermal fluctuations, but temperature here is a measure of the activity of cell extensions and thus models cell activity and motility. In accordance with previously published data suggesting that sorting might be driven by the activity of endodermal epithelial cells only (Takaku et al., 2005), we decreased the effective temperature of ectodermal cells by a factor of 2 and measured the dynamics. We found that the aggregate scale dynamics (rounding up, blob size, and boundary length) were indeed accelerated slightly by this change (Fig. S9). However, as expected, this change induced a clear difference in cell speeds between the two tissues with the endodermal cells being faster than ectodermal ones (Fig. S9). This is in direct contradiction to our experimental results from single cell tracking experiments (Fig. 3). In addition, models of cell sorting based on differential motility evolve to a final configuration in which islands of slow moving cells are surrounded by coherent streams of motile ones (Beatrici and Brunnet, 2011; McCandlish et al., 2012). The final state we obtained during cell sorting does not correspond to an internal stream of endodermal cells while the ectoderm remains passive. Indeed, we find no clear decrease in cell speeds as sorting proceeds (Fig. 3B). These results, in our opinion, clearly negate any central role for differences in cell motility in the process of cell sorting in *Hydra* aggregates.

## Discussion

### Fast dynamics of cell sorting

Our quantitative analysis of cell sorting dynamics revealed that sorting in *Hydra* aggregates is faster than published theoretical predictions. In addition, our results disagree with previous work claiming that full sorting is not observed in *Hydra* aggregates, slows down logarithmically, and could take as long as 100h (Glazier and Graner, 1993; Graner and Glazier, 1992). The largest aggregates used in our study achieved full sorting in 6-10h (Fig. 1A and Fig. S7), prior to forming a central cavity. Moreover, we only observed partial sorting on shorter time scales and for the largest aggregates studied. Quantitatively, the speed of sorting is reflected by the exponents controlling the dynamics of blob sizes and boundary lengths.

For blob size, our value of 0.49+-0.24 is higher than the result reported by (Nakajima and Ishihara, 2011) (1/3) and (Belmonte et al., 2008) (0.28). Both papers use the DAH to explain sorting in two dimensions but differ from our experiments in some key aspects. In particular, (Nakajima and Ishihara, 2011) uses periodic boundary conditions in two dimensions making any effect of the outside medium irrelevant. (Belmonte et al., 2008) use a modified Vicsek model in two dimensions in which the ectodermal cells out-number the endodermal cells three to one.

Similarly, models of sorting driven by differences in cell motility that are either intrinsic (Beatrici and Brunnet, 2011) or dependent on the cells’ local environment (Strandkvist et al., 2014) have found slower dynamics than we observed (exponents of -0.22 and -0.17, respectively). Here too, the underlying models differ from our experiments. (Beatrici and Brunnet, 2011) also used a modified Vicsek model in two dimensions in which both cell types have fixed velocities, once cell type being four times faster than the other one, which is not the case in our experimental data. Of note, they only predicted full sorting in the case where faster cells largely outnumbered slower ones and their final configuration was the opposite of what was suggested experimentally by (Jones et al., 1989). Finally, (Strandkvist et al., 2014) also use two dimensional simulations with periodic boundary conditions and tune the difference in cell motilities to be from 8-fold to 64-fold.

We attribute this difference in dynamics to the specific initial conditions used in our experiments and simulations, i.e. a three dimensional flat configuration. The equivalent configuration in two dimensions would be a thin line that, to our knowledge, has never been investigated. Since we and others (Rieu et al., 1998) have established that cell motion is mostly random during *Hydra* cell sorting, the distance that an ectodermal cell has to travel to get in contact with the outside medium is greatly reduced in a flat geometry. Of note, rounding up of the initially flat aggregates took longer than cell sorting, like in the experiments (Fig. 1A, Fig. 4A), meaning that this effect of geometry applied throughout the process. To further probe this, we ran simulations modifying the initial geometry of the aggregates to make them spherical.

This change in geometry induced only partial sorting on time scales in which similarly sized flat aggregates would fully sort. This is in agreement with previous results obtained from simulations of DAH in 2 dimensions from circular initial conditions (Graner and Glazier, 1992), the direct equivalent of the geometry tested here. Quantitatively, we found that changing the initial geometry decreased the exponent of blob size increase to 0.19 (Fig. S10), a value closer to the different theoretical predictions discussed above. Of note, these simulations also showed that the MSDs of both cell types were then diffusive (Fig. S10), further confirming that the coherent component of motion observed both in our experiments and simulations stems from the rounding up of the aggregates which occurs during sorting.

Finally, to confirm the important role of the initial geometry, we also varied the initial thickness of the square aggregates from 3 cell sizes to 7 and observed the effect of this change on sorting dynamics (Fig. S11). We indeed found that the thicker the initial aggregate, the slower the sorting further proving our hypothesis of the role of the initial geometry on sorting dynamics.

### Distinguishing between models of cell sorting

We have shown through our own simulations that DAH is sufficient to recapitulate our experimental data on both the dynamics of sorting and the behavior of single cells. Furthermore, we have incorporated differential motility into the simulations in the form of different temperatures to test whether differences in motility acted in combination with DAH to drive the sorting. Of note, it has been demonstrated experimentally that the motion of retinal cells from chick embryos during sorting was properly captured by the cellular Potts model in which temperature models membrane fluctuations (Mombach and Glazier, 1996). Varying temperature separately for both tissues is thus a proper way of modelling intrinsic differences in cell activity and motility. As a result, in this situation we observed differences in single cell behavior that we did not observe in the experimental data. Therefore, we conclude that differences in motility do not play a role in *Hydra* cell sorting.

One possibly important process that we could not test with our experimental setup is the effect of the local environment on the motility of the cells. This has been shown to suffice to drive sorting (Strandkvist et al., 2014) and has been observed experimentally (Rieu et al., 2000). Differences in motility in response to a cell’s immediate surrounding is expected from the DAH, however, and does not require intrinsic differences in motility. Indeed, in a purely endodermal cell aggregate, cell adhesions are expected to be stronger and thus cell motion to be more limited. We therefore believe that this aspect does not contradict our results.

To conclude, our multi-scale, interdisciplinary approach has answered a long-standing question regarding the mechanisms driving cell sorting in *Hydra* regeneration. We found that 1) differences in interfacial tensions between the tissues can drive sorting and 2) there are no intrinsic differences in cell motility between cell types. Our results thus rule out differential motility as a significant contributor to *Hydra* cell sorting. We confirmed these experimental results using numerical simulations. As the importance of studying physical features in the context of embryonic development is increasingly recognized, our work demonstrates that a similar approach is also fruitful in the context of regeneration, which is an exciting research area waiting to be explored in more depth with physical and biomechanical approaches.

## Materials and Methods

### *Hydra* Care

Mass cultures of the watermelon transgenic *Hydra vulgaris* line (ectoderm GFP/endoderm DsRed2), the inverse watermelon line (ectoderm DsRed2/endoderm GFP), Wnt-GFP (ectoderm DsRed expression and GFP under the control of the Wnt3 promoter (Hobmayer et al., 2000)), and AEP lines were used for experiments. AEP is the line from which embryos are obtained for making transgenic animals. Animals were kept in *Hydra* Medium (HM) composed of 1 mM CaCl2 (Spectrum Chemical, Gardena, USA), 0.1 mM MgCl2 (Fisher Scientific, Waltham, USA), 0.03 mM KNO3 (Fisher Scientific, Waltham, USA) 0.5 mM NaHCO3 (Fisher Scientific, Waltham, USA) 0.08 mM MgSO4 (Fisher Bioreagents, Pittsburgh, USA) at a pH between 7 and 7.3 at 18°C in a Panasonic incubator. Animals were cleaned daily using standard cleaning procedures from (Lenhoff and Brown, 1970). The *Hydra* were fed two to three times per week with newly hatched *Artemia* (Brine Shrimp Direct, Ogden, USA). Animals used for experiments were starved for at least 48 hours.

### Tissue Separation

The protocol is based on (Kishimoto et al., 1996) with some modifications. Animals were starved for 5-7 days before an experiment. About 10 *Hydra* were placed in 35 mm Petri dishes (CellTreat, Pepperell, USA) and cut, using sterile scalpel blades (Surgical Design Inc, Lorton, USA), below the tentacles to remove the head and above the budding zone to remove the peduncle and foot. The body columns were placed for about 2.5 minutes in ice cold HM solution, pH adjusted to 2.5 using 2 M HCl, then transferred to Dissociation Medium (DM) composed of 3.6 mM KCl (Research Products International Corp., Mt Prospect, USA), 6 mM CaCl2 (Spectrum, Gardena, USA), 1.2 mM MgSO4 (Fisher Bioreagents, Pittsburgh, USA), 6 mM sodium citrate (LabChem Inc, Zelienople, USA), 6 mM sodium pyruvate (Alfa Aesar, Ward Hill, USA), 6 mM glucose (Sigma, St. Louis, USA), 12.5 mM TES (Sigma, St. Louis, USA), 50 μg/mL rifampicin (Calbiochem, San Diego, USA), at pH 6.9 at room temperature (RT) following (Flick and Bode, 1983). The dishes containing body columns in DM were taped and swirled to promote separation of tissues. Success rate was low with only around 10% of body columns fully separating and around 20% showing partial separation. In this latter case, ectoderm and endoderm pieces would be manually cut free. After separation, samples were further cut with scalpel blades to yield pure pieces of either tissue type.

### Cell aggregates

Aggregates were prepared according to (Gierer et al., 1972), with some modifications. About 100 *Hydra* body columns from various strains were prepared by cutting off the head and peduncle/foot and washed 3 times with DM before 1 hour incubation in DM on ice. The body columns were mechanically dissociated into a single cell suspension with vigorous trituration. The cells were centrifuged in an Allegra X-15R Centrifuge (Beckman Coulter, Brea, USA) at 4C, 200g, for 5 minutes and washed twice with ice cold DM. About 1 mL of cell suspension was made from 100 body columns by washing the pellet of single cells through a Flacon nylon 40 μm nylon mesh filter (Corning Incorporated, Corning, USA). 100 μL aliquots of this cell mix were placed in separate wells of 96 well V-shaped plates (Nunc, Roskilde, Denmark) and the plate was centrifuged at 1,000g for 5 min. Aggregates were cultured in 100% DM for the first 4 hours after which 100 μL of HM was added to each well leading to a 50/50 mix of HM and DM. At 24 hours, the aggregates were transferred using glass Pasteur pipets to a solution of 70:30 DM:HM. Finally, at 48h, they were transferred to 100% HM for the rest of regeneration. Throughout, aggregates were kept at 18°C. Imaging started 1 hour after the aggregates were made by carefully transferring them to 96 well flat-bottom plates (Nunc, Roskilde, Denmark). The aggregates were imaged every 2 to 5 minutes using GFP and RFP channels and 3 planes z-stack on an Olympus IX81 inverted microscope (Olympus Corporation, Tokyo, Japan) using an ORCA-ER camera (Hamamatsu Photonics, Hamamatsu, Japan) and Slidebook software (version 5, Intelligent Imaging Innovations, Inc, Denver, USA).

By visually controlling cell density in the cell mix prior to centrifugation, we prepared aggregates of different initial sizes, ranging from 10^2^ to 10^4^ cells. Of note, only the largest of these (roughly over 10^3^ cells) fully regenerate and we thus focused on these in our experimental analysis.

### Analysis of sorting dynamics

Two channel z-stack images of aggregates were used for analysis to determine boundary ratio and blob size measurements. The z-stacks were first converted to maximum intensity projection RGB image sequences in ImageJ (http://imagej.nih.gov/ij/, NIH, Bethesda, USA). Using a semiautonomous MATLAB (MathWorks, Natick, USA) script, the red and green channels were normalized to each other based on each channel’s average intensity. The normalized image was then segmented into three regions—background, ectoderm, and endoderm—using the function kmeans (Fig. 1B). We measured the ‘length’ of the boundary between the endoderm and the ectoderm by taking the sum of all the points between the ectoderm and endoderm segments. The boundary ratio was defined as this length divided by the perimeter of the aggregate. Of note, we find this measurement to be directly proportional to the sorting index used in other papers. This index is calculated as the fraction of neighboring cells which are of the opposite cell type, averaged over all cells. On average, each cell will thus have a boundary length equal to the mean sorting index times its contour length. Assuming that all cells have similar sizes, the total boundary length will then be the average boundary length per cell times the number of cells which is thus also proportional to the sorting index.

For blob size, we calculated the segmented image’s 2-D Fourier transform 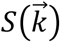 and the typical blob size as:

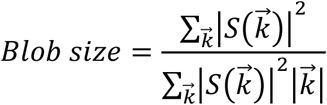

The blob size and boundary ratios from 17 aggregates (5 technical replicates) were individually linear fitted on a log-log plot to determine the power law exponent. The blob size and boundary exponents were averaged over these 17 measurements and their standard deviations calculated. For plotting purposes, blob sizes and times were normalized by dividing them by their initial value. Of note, this normalization doesn’t alter the exponents that were obtained.

To estimate volumes from two-dimensional imaging, the aggregates were assumed to be ellipsoidal. We fit an ellipse over the segmented image in ImageJ and used its minor axis length for the width and girth and the major axis as the length of the ellipsoid.

The sorting time, the time it takes for the ectoderm and endoderm to be completely separated (no ectoderm cells remain inside the endoderm), was found by having two people watch the sorting videos and manually record the time points when the aggregates become sorted. We then averaged and calculated the standard deviation of the manually recorded sorting times.

### Micropipet aspiration

Tissue pieces were used right after cutting following tissue separation. Glass capillaries (model 1B100F-4, World Precision Instruments, Sarasota, USA) were pulled into micro-pipettes using a horizontal laser based micropipette puller P-2000 (Sutter Instrument, Novato, USA). The resulting needles were manually cut to yield an opening of approximately 50 microns (smaller than pieces of interest but larger than one cell). Before the experiments, they were treated with Sigmacote (Sigma, St. Louis, USA) following the manufacturer’s protocol to make them non-adhesive. They were then mounted onto a needle holder attached to an M-152 micromanipulator (Narishige USA, Amityville, USA). The assembly was connected by hermetically sealed tubing to a plastic syringe used as a water reservoir and mounted onto a stand allowing for manual variation of the syringe’s height. Using the micromanipulator, the needle tip was put in contact with the piece of interest before lowering the syringe’s level to apply negative pressure. The aspiration of the piece was imaged every 5-10s under a MZ16FA stereo microscope (Leica Microsystems, Wetzlar, Germany), using a SPOT RT3 camera (Model 25.1, Diagnostics Instruments, Sterling Heights, MI, USA) controlled by SPOT Basic 5.1 software (Diagnostic Instruments). For surface tension estimates, the radius of the piece was measured from the images in ImageJ as the square root of the projected area divided by π. The radius of the needle was also measured in ImageJ. The height difference between the water reservoir and the Petri dish containing the sample was slowly manually increased until aspiration began. The height difference was recorded at that time and translated into a critical pressure value according to:

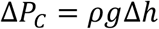

Where Δ*P_C_* is the critical pressure, *ρ* is the density of the medium and Δ*h* is the height difference between the water reservoir and the Petri dish.

This led to a surface tension estimate per piece. The values presented in the results section are averaged from 13 independent endoderm and 14 independent ectoderm pieces.

For viscosity estimates, the retracted length as a function of time was manually measured in ImageJ and fit in MATLAB as an exponential function of the form: 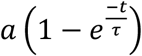. *a* and *τ* were fit parameters with:

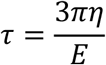

Where *E* and *η* are the elastic modulus and viscosities, respectively. This led to an estimate of the viscosity for each tissue piece. The values presented in the results section are averaged from 10 independent endoderm and 9 independent ectoderm pieces.

### Fusion

The two tissue layers obtained from tissue separation were cut into smaller pieces. Within 5 minutes post-cutting, the two tissue pieces of interest were either put into contact using the hanging drop technique described in (Schoetz, 2008) or manually brought in contact using hair pins. The fusion process was imaged every minute either on an Olympus IX81 inverted microscope or using an EVOS FL Auto Cell Imaging System (Thermo Fisher Scientific, Waltham, USA).

### Rounding up

Pieces from either tissue were manually cut with sterile razor blades into oblong shapes in DM and imaged every minute for 2-3h under a Leica MZ16FA stereo microscope. Using ImageJ, each piece was fitted by an ellipse at each time point and the aspect ratio was computed as the ratio of the long axis to the short one as a function of time. The dynamics was then fit in MATLAB by an exponential decay function of the form

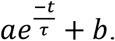

*a*, *b* and *τ* were fit parameters and the characteristic time *τ* was taken to be 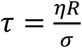 with *η* the viscosity of the tissue, *σ* its surface tension and *R* the radius of the piece of interest (Mombach et al., 2005). Radii were measured as the geometric mean of the axes of the fitting ellipse at the final time point. Using viscosity estimates from micro-pipet aspiration experiments, we measured the surface tension of each piece this way and the results presented are averaged over 17 independent endoderm and 15 independent ectoderm pieces.

### Single cell dynamics

Aggregates containing 5% of their cells from watermelon animals and 95% from AEP animals were prepared as described above. After 1h in DM, they were imaged on a Scientifica multiphoton imaging setup (Scientifica, Uckfield, UK) coupled to a MaiTai ultrafast laser (Spectra Physics, Santa Clara, CA, USA) set to 980 nm through a 20x water immersion XLUMP PlanFL objective (Olympus Corporation, Tokyo, Japan). Optical slices in both the RFP and GFP channels were acquired at 3 microns-slices and averaged over 6 acquisitions. This procedure was repeated every 5 to 10 minutes over 4-6h.

Single cells were detected separately for each tissue using the *3d object counter* plugin in ImageJ and their center of mass at each time point recorded. This data was analyzed in MATLAB by reconstructing single cell trajectories by usual tracking algorithms freely available online (http://www.mathworks.com/matlabcentral/fileexchange/42573-particle-point-analysis?focused=3791012&tab=function). From these tracks, we calculated mean square displacements, speeds, auto-correlation functions and directionality according to the usual definitions. Values reported here are averaged over all trajectories for each cell type. The data presented in the results section are from one representative experiment out of eight.

For center of mass corrections, we used mosaic aggregates containing 5% of cells from HyWnt3 promoter::GFP animals allowing us to track ectodermal cells in the RFP channel and 95% from AEP animals. During the 1h period in DM, aggregates were stained with a Syto12 nuclear dye (Thermo Fisher Scientific, Waltham, USA) diluted to 1:500 in DM. Aggregates were washed twice in DM before imaging on the 2-photon microscope. The analysis was performed in the same way as above. Center of mass was calculated from the mean position of all detected nuclei at each time step and the center of mass position was subtracted from both nuclei and ectodermal cell positions. The corrected positions were then used to calculate mean square displacements.

### Numerical simulations

We used the freely available CompuCell3d (Swat et al., 2012) software to perform simulations of a Cellular Potts model. Details on how these simulations are performed can be found in the software’s manual (available at: http://www.compucell3d.org/Manuals). In our case, we used a Hamiltonian *H* composed of three contributions. The first one modelled cell-cell contacts and was written as follows:

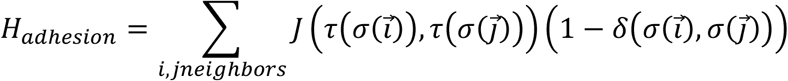

Where the summation applies over all pairs of adjacent pixels 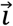 and 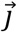, 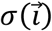 is the ID number of the cell occupying the pixel 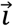 and 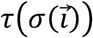 the identity (endoderm, ectoderm, or medium) of that cell and *δ*(*x*, *y*) is the Kronecker function. This formulation means that energy only applies to neighboring pixels that belong to different cells and the energetic cost of that adhesion depends on the identity of both cells involved. For the adhesion energies, we used the following parameters:

| Tissues | $J$ |
| --- | --- |
| Endoderm/medium | 300 |
| Ectoderm/medium | 200 |
| Endoderm/endoderm | 2 |
| Endoderm/ectoderm | 75 |
| Ectoderm/ectoderm | 100 |

The relative values of these parameters were chosen to fulfill the following criteria: the adhesion energy between the two tissues has to be intermediate between the two homotypic adhesion energies to account for complete engulfment; the effective surface tension of the endoderm has to be double that of the ectoderm, in agreement with experiments. The surface tension of one tissue was estimated as:

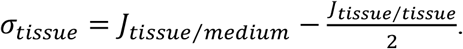

The second contribution to the Hamiltonian was the limited compressibility of cells which means that they resist any deviation from a target volume leading to the following formulation:

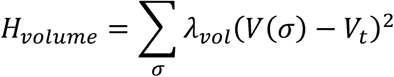

Where the summation applies over all cells, *λ_vol_* is the inverse compressibility of the cells, *V*(*σ*) is the volume of cell *σ* and *V_t_* is the target volume. In our simulations, we used the same compressibility for both cell types with the following parameters *λ_vol_* = 20 and *V_t_* = 64.

The last contribution represents the cell’s membrane tension. Numerically, this means that there is an energy penalty for the surface of each cell if it deviates from a target value, leading to the following formulation:

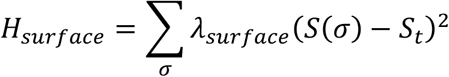

Where the summation applies over all cells, *λ_surface_* is the inverse membrane compressibility of the cells, *S*(*σ*) is the surface of cell *σ* and *S_t_* is the target surface. In our simulations, we used the same membrane compressibility for both cell types, with the following parameters *λ_surface_* = 5 and *S_t_* 96.

Overall, the Hamiltonian controlling the dynamics was the sum of the three contributions above. The activity of the cells was then represented as a temperature parameter that allows them to overcome local energy barriers to reach the energetically optimal situation. In our simulations, the temperature was set to 1000 with the exception of our simulations of differential motility that kept the same temperature for the endoderm but used 500 for the ectoderm.

Simulations were initialized, in most cases, as rectangular random mixtures of cells (initialized as cubes with 4 pixel sides) from both cell types. The width and length of the rectangles were kept equal and were varied from 30 pixels to 140 pixels while the thickness was kept at 12 pixels, except for the data presented in Fig. S11. For sorting with spherical initial conditions, we initiated the simulation as a random mixture of both cell types in a sphere with a radius of 40 pixels. To simulate fusion, two spheres, in contact, for a single tissue type were initiated with a radius of 22 pixels each.

For analysis, data were saved at intervals ranging from 2 to 30 simulation steps depending on the system size. An image representing the horizontal slice in the middle of the aggregate’s height was recorded and detailed data on the identity of each pixel were saved. The horizontal slices were used to calculate blob sizes and boundary lengths in the same way as described in the *Sorting dynamics* section. The saved time series data were used to reconstruct the three-dimensional dynamics of each cell’s center of mass. For cell sorting, we ran triplicates of the simulations at ten different sizes. For each size, one exponent for blob size and one for boundary length were obtained by linear fitting their dynamics on a log-log plot in MATLAB. The values reported are means of the values obtained for the seven largest aggregate sizes leading to a total of 21 simulations. For single cell dynamics, the data presented come from a single simulation at the largest size studied, but which is representative of all simulations.

### Immunohistochemistry

Aggregates from AEP animals were prepared as described above and fixed at different time points in 4% paraformaldehyde in HM for 15 min at RT. They were washed 3x10 min in phosphate buffered saline (PBS) and permeabilized for 5min in PBS supplemented with 0.5% Triton X-100 (Sigma, St. Louis, USA). A blocking solution was prepared using 1% bovine serum albumin (BSA) (Fisher Bioreagents, Pittsburgh, USA), 10% fetal bovine serum (Thermo Fisher Scientific, Waltham, USA) and 0.1% Triton X-100 in PBS. Aggregates were blocked in that solution for 2h at RT. The samples were incubated in primary anti-*Hydra* laminin, mAb 52 antibody (Sarras et al., 1994), diluted to 1:200 in the blocking solution, overnight at 4°C. Negative controls were performed by omitting the primary antibody and using blocking solution alone. Next, samples were washed 6-8 times in PBS over the course of 3-5 hour at RT before incubating overnight in a 1:500 dilution, in blocking solution, of an anti-mouse HRP secondary antibody (Enzo Life Sciences, Farmingdale, USA). The next day, samples were again extensively washed in PBS before a 1h incubation in PBT (1:2000 Tween (Sigma, St. Louis, USA) and 0.2% BSA in PBS). Detection of the HRP secondary antibody was performed in PBT supplemented with 1:10000 H2O2 (Avantor, Central Valley, USA) and 1:1000 NHS-fluorescein (Thermo Fisher Scientific, Waltham, USA) for 30 min. Samples were washed multiple times at RT and overnight at 4°C in PBS. Samples were imaged on an Olympus IX81 inverted microscope. The resulting images were analyzed by measuring the averaged signal intensity in the middle of the aggregate. This value was normalized by the same measurement performed on the negative controls. Results from four different experiments were averaged and their standard deviations calculated.

### Centripetal aggregations

Endoderm and ectoderm tissue pieces were dissociated into single cell suspensions in the same way as in preparing cell aggregates. Cell concentrations were measured using a Brightline hemacytometer (Sigma, St. Louis, USA) and equalized by adding DM to the most concentrated cell suspension. 800μL of these suspensions were placed in wells of a 24well plate (Genesee Scientific, San Diego, USA) and placed on a DS-500E rotary shaker (VWR International, Radnor, USA) for 30min at 75rpm. The resulting aggregates were then imaged under a Leica MZ16FA stereo microscope. Images were analyzed in ImageJ to extract the projected area of each resulting aggregate.

## Acknowledgments

We thank E. Cary and G. Lee for *Hydra* care and tissue separation, R. Merks for his help with numerical simulations, W.-J. Rappel, T. Goel and P. Diamond for comments on the manuscript, and X. Zhang for providing the mAb52 antibody.

